# Diurnal Chorusing in Nine Species of North American Frogs

**DOI:** 10.1101/336149

**Authors:** Malcolm L. McCallum, Jamie L. McCallum

## Abstract

A year-long survey of survey of diurnal frog calling behavior was conducted at the Red River Research and Education Park in Shreveport (Caddo Parish), Louisiana to investigate the prevalence of daytime breeding choruses in the species present at the park. We determined that 60% (9/15) of species known from the area participated in daytime breeding choruses. Four of these were new species reports for the behavior. One species was identified entirely based on a daytime chorus from outside the normal breeding season. Although we believe daytime chorusing is widespread in frogs, several species did not diurnally chorus during our study. Diurnal calling may be an important indicator of the peak breeding season for a species and it may also be a useful tool at times when night surveys are impossible.

## Introduction

Much research has focused on calling behavior in frogs (Gerhardt 1994), but these are typically focused at night. In fact, virtually all descriptions of anuran calling behavior are based on nocturnal surveys and observations. However, many species continue calling for mates throughout daytime hours (pers. obs.), despite the lack of attention to this behavior. Although there are mentions of diurnal breeding choruses in the literature (e.g. Cane Toad [*Rhinella marina*] [Krakauer 1968; Meshaka et al. 2011], the Coqui [*Eleutherodactylus coqui*] [Meshaka et al. 2011], Greenhouse Frog [*Eleutherodactylus planirostris*] [Goin 1947; Meshaka et al. 2004], and the Cuban Treefrog [*Osteopilus septentrionalis*] [Meshaka et al. 2011]), few studies specifically investigate the prevalence of diurnal breeding choruses in frogs. One conducted at the U.S. Department of Energy’s Savannah River Site in South Carolina, summer 1997 (Bridges and Dorcas 2000) used automated recording systems to continuously record calling behavior. It found that *Acris gryllus, Lithobates catesbieanus, L. clamitans* and *Gastrophryne carolinensis* regularly called during the day, albeit at lower levels than during the night. *Lithobates sphenocephalus, Hyla cinerea, H. femoris*, and *H. chrysoscelis* day-called sporadically, and *H. gratiosa* did not call during the day. Spring Peepers (*Pseudacris crucifer*) and Blanchard’s Cricket Frog (*Acris blanchardi*) oviposit during the day and night in New York (Wright 1914) and diurnally chorus early in the season (Kenney and Stearns 2015). Most studies of frog calling ignore daylight hours, and many guidelines (e.g. NAAMP, Frogwatch) recommend surveys in the early evening hours. Hence, the behavior appears otherwise undocumented. There has been much discussion about the importance of natural history in the face of conservation needs (Bury 2006, McCallum and McCallum 2006), and diurnal chorusing is certainly an area in which our knowledge is lacking.

## Materials and Methods

The study site was an oxbow lake and wetland at the Red River Research and Education Park (a.k.a. C. Bickham-Dickson Park) in Shreveport (Caddo Parish), Louisiana (Population = 400,000). This 249 ha urban wetland (Fig. 1) surrounds an oxbow lake that is connected by a small channel to the Red River during most of the year. During the winter, the Red River frequently inundates the park. The vegetation in the park is a mix of native and exotic species (MacRoberts et al. 2008).

**Figure 1.**
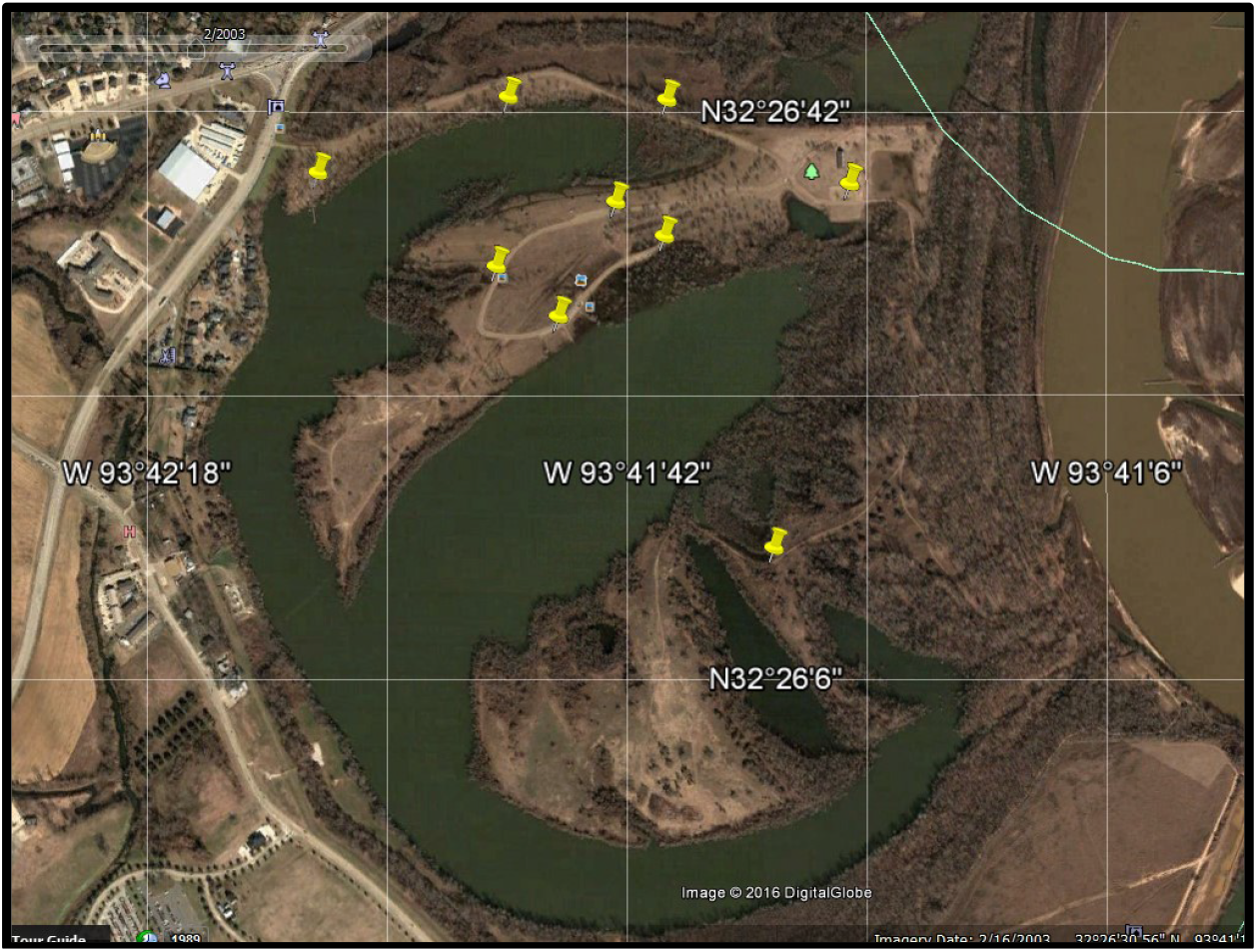
Historical aerial imagery of the Red River Research and Education Park (Shreveport [Caddo Parish], Louisiana] in 2003-2004. Yellow markers indicate each of nine observation stations (Source: Google Earth).

We visited the Red River Watershed Research and Education Park between 1100 and 1300 hrs on nearly a daily basis from 20 September 2003 through 4 January 2005, totaling 249 visits. Some stations could not be visited during floods. Each visit lasted 60 – 120 min, depending on the amount of avian activity present on a given day (this investigation was coupled with an avian survey). We drove the perimeter road with the windows down and stopped at nine watch stations (Fig. 1, 2). Whenever we noted frogs calling, we stopped and left the vehicle, listened quietly, then logged the location and species observed in a notebook. All sessions were recorded with a hand-held digital audio recorder for later review and verification. For the purpose of this study, isolated single calling males were excluded because these were more characteristic of a rain call than calling for mates. We also recorded ambient temperature and wind speed using a Kestrel^®^ hand-held weather unit and then noted any precipitation during each stop. We could not survey frog calling at night because of city ordinance, locking of the park gate outside of working hours, and a lack of funds for automated recorders. The minimum number of stops during each visit was one per each of the nine stations selected at the beginning of the study. Results were statistically analyzed with regressions using MiniTab 13.0.

**Figure 2.**
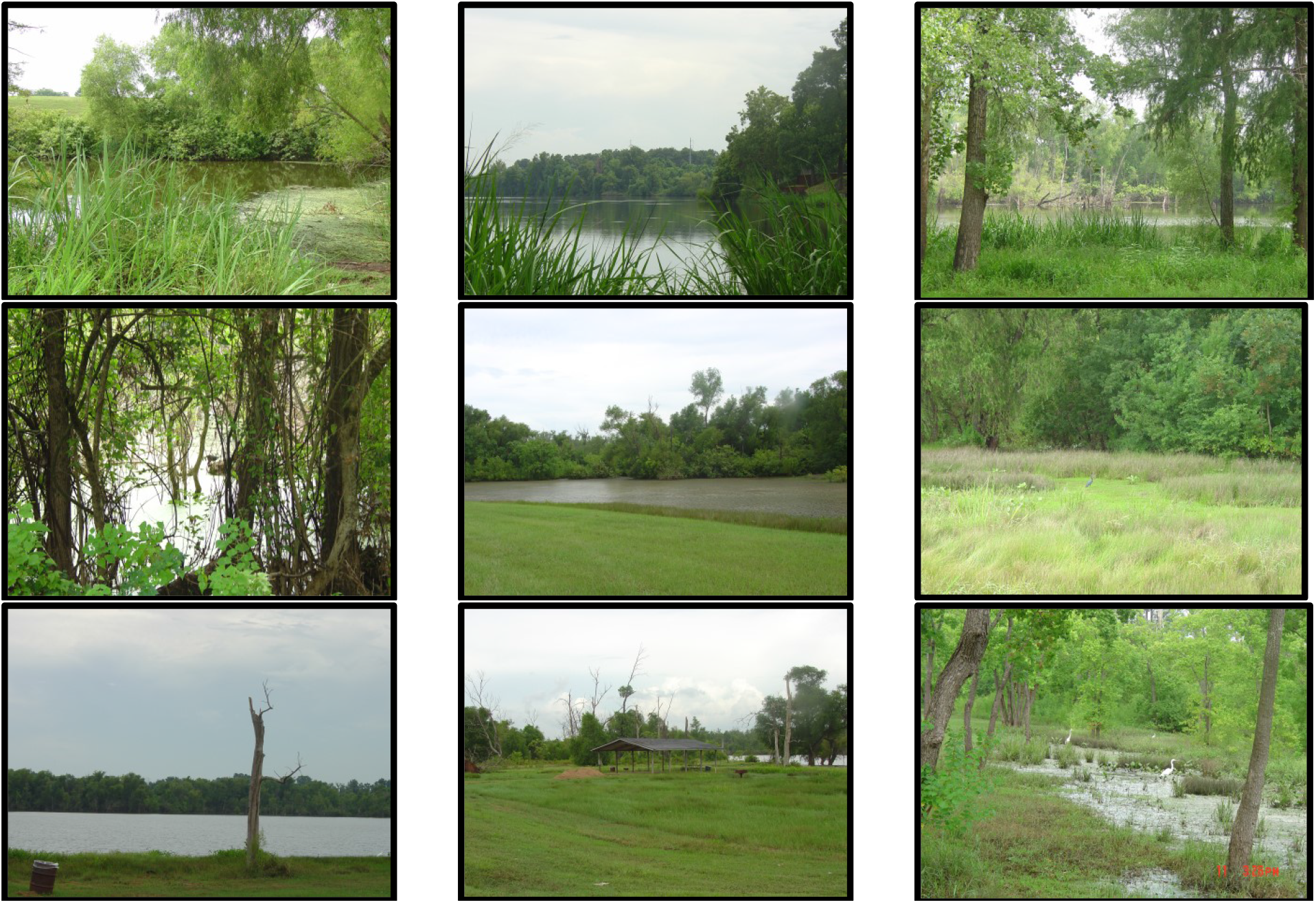
Habitats at nine listening stations at the Red River Research and Education Park (Shreveport [Caddo Parish], Louisiana) in 2003 – 2004. (Photos by Jamie McCallum).

## Results

Nine species of anurans called diurnally during our study (Table 1). Among those, only the American Toad (*Anaxyrus americanus*) had not been physically observed at the park or in the immediate surrounding area. The Pickerel Frog (*Lithobates palustris*) and Cope’s Treefrog (*Hyla chrysoscelis*) were known from the surrounding area, were not previously observed at the site, but detected via daytime choruses. The Green Treefrog (*H. cinerea*), Gray Treefrog (*H. versicolor*), and Cajun Chorus Frogs (*Pseudacris fouquettei*) were previously recorded at the park, but did not call during our daily visits.

**Table 1.** Observations of Anuran species at the Red River Research and Education Park, Shreveport, Louisiana and records of diurnal breeding choruses.

| Species | Present?* | Calling Season (NAAMP) | Detection Window |  | Detectability |  |
| --- | --- | --- | --- | --- | --- | --- |
|  |  |  | Earliest Diurnal chorusing | Latest Diurnal chorusing | No. of visits from first to last calling day N/T (%) | Total Visits N/T (%) |
| Blanchard's Cricket Frog (*Acris blanchardi*) | Yes | March – Oct.** | 15 March 2004 | 3 September 2004 | 72/87 (83%) | 72/248 (29%) |
| Eastern Narrowmouth Toad (*Gastrophryne carolinensis*) | Yes | May – July | 4 June 2004 | 11 July 2004 | 9/19 (47%)<br>[28 June – 1 July = 4/9 (44%) of calling days] | 9/248 (3.6%) |
| Fowler's Toad (*Anaxyrus fowleri*) | Yes | April – July | 4 June 2004 | 1 July 2004 | 8/15 (53%)<br>[27 June – 1 July = 5/8 (63%) of calling days] | 8/248 (3.2%) |
| American Bullfrog (*Lithobates catesbeianus*) | Yes | April – July | 28 March 2004 | 27 June 2004 | 8/49 (16%) | 8/248 (3.2%) |
| Southern Leopard Frog (*Lithobates sphenoccephala*) | Yes | January – July | 25 January 2004 | 1 March 2004 | 5/30 (17%) | 5/248 (2%) |
| American Toad (*Anaxyrus americanus*) | No | Est. Mar – June*** | 27 June 2004 | 19 July 2004 | 2/12 (17%) | 2/248 (0.8%) |
| Bronze Frog (*Lithobates clamitans*) | Yes | March – July | 27 June 2004 | 11 July 2004 | 2/8 (25%) | 2/248 (0.8%) |
| Pickereel Frog (*Lithobates paulustrus*) | Maybe | March | 1 February 2004 | 1 February 2004 | -- | 1/248 (0.4%) |
| Cope's Treefrog (*Hyla chrysoscelis*) | Maybe | March – July | 17 October 2004 | 17 October 2004 | -- | 1/248 (0.4%) |
| Bird-voiced Treefrog (*Hyla avivoca*) | Maybe | April – July | -- | -- | -- | 0/248 (0%) |
| Green Treefrog (*Hyla cinerea*) | Yes | March – July | -- | -- | -- | 0/248 (0%) |
| Gray Treefrog (*Hyla versicolor*) | Yes | April – June | -- | -- | -- | 0/248 (0%) |
| Squirrel Treefrog (*Hyla squirrella*) | Maybe | June | -- | -- | -- | 0/248 (0%) |
| Spring Peeper (*Pseudacris crucifer*) | Maybe | January – May | -- | -- | -- | 0/248 (0%) |
| Cajun Chorus Frog (*Pseudacris fouquettei*) | Yes | December – May | -- | -- | -- | 0/248 (0%) |
| Rio Grande Chirping Frog (*Eleutherodactylus cystignathoides*) | Maybe | ? | -- | -- | -- | 0/248 (0%) |

\*Yes = physically observed in park, No = not physically observed at park or in area, Maybe = not physically observed in park but present nearby.
\*\* NAAMP surveys suggest March – July, but our personal observations in the area suggest this frog calls through October at night.
\*\*\*No NAAMP records in northern Louisiana, southern Arkansas or northeastern Texas. These dates based on the closest NAAMP route at Vicksburg National Battlefield, Mississippi (510610).

The detection window while using diurnal chorusing was similar to use of nocturnal choruses for two species (Blanchard’s Cricket Frog and the Pickerel Frog). Six of the species had daytime detection windows that were shorter than the known night chorus (Table 1). Cope’s Treefrog breeds from March – July, but was only observed chorusing diurnally in October (Table 1).

There was sufficient data to assess interactions among ambient temperature, wind speed and chorusing in five of the nine species observed calling diurnally (Table 2). Among these five species that diurnally chorused, only Blanchard’s Cricket Frog responded to ambient temperature or wind speed (*r^2^* = 0.307). Depending on the date, temperature and wind speed influenced if this species diurnal chorused (Table 3).

**Table 2.**
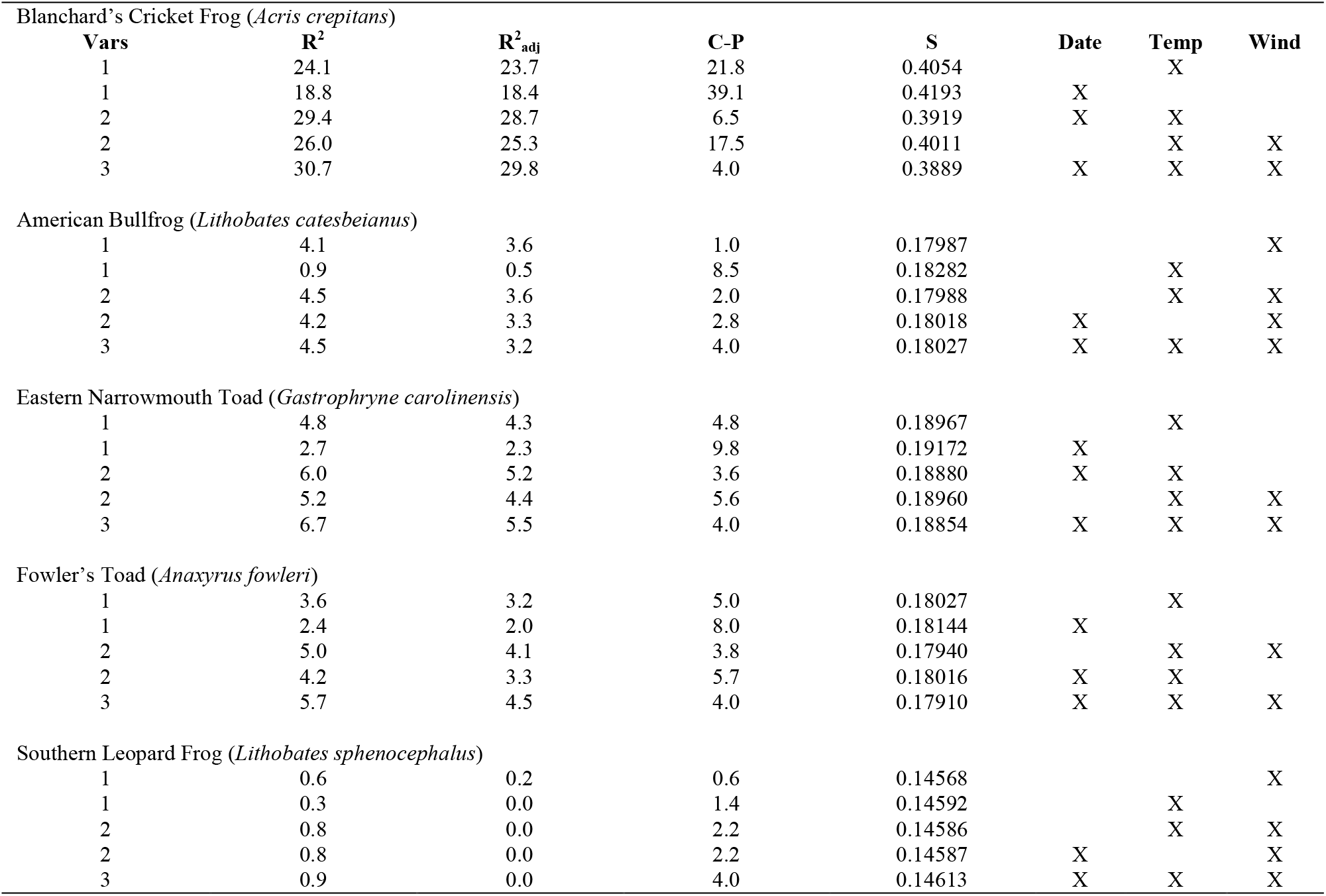
Results of Best Subsets Regression results for the possible interaction between the date, temperature, and wind speed on expression of diurnal calling in fives species of frogs.

**Table 3.** Results from Binomial logistic regression for the influence of the date, wind speed and temperature on the occurrence of calling by male Blanchard’s Cricket Frogs (*Acris blanchardi*).

| **Response Information** |  |  |  |  |  |  |  |
| --- | --- | --- | --- | --- | --- | --- | --- |
| **Variable** | **Value** | **Count** |  |  |  |  |  |
| Calling | 1 | 72 |  |  |  |  |  |
| No calling | 0 | 159 |  |  |  |  |  |
| Missing values | -- | 17 |  |  |  |  |  |
| **Logistic Regression Table** |  |  |  |  |  |  |  |
| **Predictor** | **Coefficient** | **SE Coefficient** | **Z** | **P** | **Odds Ratio** | **95% CI** |  |
| Constant | -191.41 | 62.32 | -3.20 | 0.001 |  |  |  |
| Date | 0.00509 | 0.001641 | 3.10 | 0.002 | 1.01 | 1.00 | 1.01 |
| Temperature | 0.16651 | 0.03313 | 5.03 | < 0.001 | 1.18 | 1.11 | 1.26 |
| Wind Speed | 0.3466 | 0.1440 | 2.41 | 0.016 | 1.41 | 1.07 | 1.88 |
| Log-Likelihood | -101.431 |  |  |  |  |  |  |
| **Test that all slopes are zero** |  |  |  |  |  |  |  |
| **G** | **df** | **P** |  |  |  |  |  |
| 83.784 | 3 | < 0.001 |  |  |  |  |  |
| **Goodness-of-Fit Tests** |  |  |  |  |  |  |  |
| **Method** | **Chi Square** | **df** | **P** |  |  |  |  |
| Pearson | 181.425 | 227 | 0.988 |  |  |  |  |
| Deviance | 202.862 | 227 | 0.874 |  |  |  |  |
| Hosmer-Lemeshow | 26.494 | 8 | 0.001 |  |  |  |  |

## Discussion

Previous observations suggest that using a limited listening window in the evening may cause some species to go undetected (Bridges and Dorcas 2000). In fact, our data support this concern. Cope’s Treefrog was not previously observed at the park. However, we detected it chorusing in the fall. This species would have gone undetected had we not surveyed the entire year. Whether fall diurnal chorusing was errant or typical behavior for the region is not definite. However, there have been observations of overwintering Cope’s Treefrog tadpoles in Shreveport, (McCallum and McCallum 2004) of a size suggesting fall oviposition. If this species breeds in the fall, its tadpoles would need to overwinter before metamorphosing.

Blanchard’s Cricket frog breeding choruses take place between March – October in the Arkansas Ozarks (McCallum 2003; Trauth et al. 2004). Females with large vitellegenic ova are present from April – August and males have sperm present throughout the year in most of Arkansas (McCallum et al. 2011). Day-calling is prominent from March – September in northwestern Louisiana. By September, females with yolked egg clutches are rare in Arkansas and the population has largely turned over to young-of-the-year (McCallum 2003; McCallum et al. 2011). Considering the latitudinal differences between northwestern Louisiana and most of Arkansas, day-calling appears to line up very closely with the presence of ripe females in the population. We pose that day-calling may indicate the peak breeding activity, and potentially reflect testosterone levels in male frogs. However, more in-depth study is needed to validate this hypothesis.

Our study suggests that diurnal chorusing by male frogs might be more widespread than previously known and that failure to consider this may result in undetected but present species in status surveys and inventories. We suspect strongly that this behavior is much more common across species than previous reports would suggest. We found four species of frogs that had not previously been reported in the peer-reviewed literature to chorus during the day (e.g. American Toad, Fowler’s Toad, Blanchard’s Cricket Frog, and the Pickerel Frog). This may constitute an important tool and consideration for both applied and theoretically-focused herpetologists.

